# A Quadrivalent mRNA immunization elicits potent immune responses against vaccinia and monkeypox viral antigens – a step closer to a broad orthopoxvirus vaccine

**DOI:** 10.1101/2023.04.23.537951

**Authors:** Caixia Su, Yang Wen, Xiya Geng, Chunmei Yang, Quanyi Yin, Yelin Xiong, Zhihua Liu

## Abstract

The global outbreak of the 2022 monkeypox virus infection of human raised the public health concerns of the threat of human-to-human transmission of zoonotic diseases. Given the evidence that other orthopoxviruses including cowpox and camelpox were also reported infectious to human, and that the reemerging risk of smallpox as a bioterrorist or accidental laboratory escape exists, there is an urgent need to develop a poxvirus vaccine with a broad protection of orthopoxviruses to stockpile for future emergency. Extensive studies of vaccinia virus (VACV) suggested that multiple VACV antigens, such as A27, L1, A33 and B5, showed high level similarity in terms of immunogenicity to their respective homologous antigens of other orthopoxviruses. These findings paved the ground for VACV antigens to be used as potential vaccine targets for development of a universal poxvirus vaccine. In this study, we construct a novel poxvirus vaccine candidate, mRNA-ALAB-LNP, encoding four vaccinia viral antigens A27, L1, A33 and B5. Strong anti-L1-specific antibody and moderate anti-A33-, anti-A27- and anti-B5-specific antibody responses were induced in mice after a single immunization. The antibody responses to all four antigens were significantly boosted after the second shot with all IgG titers >5 logs and highest being anti-A33 IgG. The high level of binding antibodies showed potent neutralizing capability against vaccinia virus. Specific IFN-γ responses were detected to all four antigens with the highest cellular response being that induced by the same antigen, A33. When evaluating the cross reactivity, equivalent or better serum IgG responses were seen in responses to corresponding monkeypox antigens A35, M1, A29 and B6, in comparison to vaccinia antigens. Apparently, the mRNA vaccine encoding four vaccinia antigens induced immunity not only to vaccinia virus but also to monkeypox, suggesting that the mRNA-ALAB may be a candidate for potential vaccine development against infection of monkeypox, smallpox and possibly other orthopoxviruses.

## Introduction

Orthopoxviruses belong to the family Poxviridae and the genus orthopoxvirus contain 12 species including smallpox, monkeypox, vaccinia, camelpox and cowpox virus. Smallpox (caused by variola virus) was one of the most devastating human diseases that caused millions of deaths before it was eradicated in 1980 (Derrick B.1988). Following the eradication of the naturally-occurring smallpox, risk of variola infection is minimum with the possible sources of infection being deliberate release of virus for military purposes or accidental escape of virus from a laboratory. During 2021 to 2022, monkeypox outbroke in African countries with 1329 cases and 68 deaths reported followed by the first case outside Africa spotted in the UK and subsequently the virus spread within 6 months globally with total cases of over 80 thousands (Mitjà et al., 2023). The sequence analysis showed that the virus belongs to the West African branch (Mitjà et al., 2023), one of the two distinct clades of monkeypox, despite that the understanding of the factors leading to the current outbreak is limited (Gessain et al., 2022; Lum et al., 2022). The monkeypox epidemic indicated that monkeypox virus, as a pathogenic orthopoxvirus, can become another pathogen potentially threatening public health and safety after smallpox.

To our knowledge, monkeypox is not the only orthopoxvirus that possess potential threats. Cowpox infection of human occurred in Europe and adjacent Russian states. One common host is the domestic cat, from which human infections are most often acquired (Bennett et al., 1990). Cowpox virus has also infected a variety of animals in European zoos, such as elephants, resulting in human infection (Kurth et al., 2008). Camelpox is another orthopoxvirus that is very closely related to the variola virus and vaccinia. The camelpox virus most often affects members of family Camelidae. However, recent studies show that the disease can be transmitted to both humans and arthropods (Duraffour et al., 2011; Jezek et al., 1983). The most recently described orthopoxvirus species that infected human is the Alaskapox virus, first isolated in 2015 (Gigante et al., 2019). At this point, in addition to risk of deliberate military release and accidental laboratory release of smallpox, there is a threat of zoonotic orthopoxvirus to human health. Development of a more effective poxvirus vaccine with a broader protection and scalable manufacturing process is imminent.

To develop an effective broad orthopoxvirus vaccine, understanding of the variations between family members is important. The genome of vaccinia (Goebel et al., 1990), variola virus (Massung et al., 1994), monkeypox (Shchelkunov et al., 2001), camelpox (Afonso et al., 2002), ectromelia virus (Mavian et al., 2014) and cowpox virus (Shchelkunov et al., 1998) have been sequenced and the results demonstrated that all viruses are morphologically indistinguishable and antigenically related. Any prior infection with one virus will provide some protection against each other members of the genus (Fenner et al., 1989). Using a poxvirus-specific tool, accurate gene sets for viruses with completely sequenced genomes in Orthopoxvirus were predicted, such that, in all existing Orthopoxvirus species, no individual species has acquired protein-coding genes unique to that species (Hendrickson et al., 2010). This was the foundation for cowpox virus and vaccinia virus being effective vaccines against smallpox and monkeypox. Based on the above, we reasonably speculate that a vaccine targeting proper vaccinia viral antigens may provide protective immune responses against broad orthopoxviruses.

The vaccinia virus, like other orthopoxviruses, contains a linear double-stranded DNA genome (Gessain et al., 2022), encoding about 200 proteins, including various proteins and enzymes required for virus replication, virus assembly, host restriction, pathogenicity and other processes. The virus often exists in two different infectious forms, the intracellular mature viruses (IMV) and extracellular enveloped viruses (EEV), whose surface glycoproteins infect cells using different mechanisms (Franceschi et al., 2015). Early studies show that the four membrane proteins A27, L1, A33, and B5 of vaccinia virus are involved in the adsorption, binding, and intercellular transmission of virus-infected cells (Chung et al., 1998; Foo et al., 2009; Smith et al., 2002). The A27 protein and the L1 protein on IMV are generally considered to mediate the attachment and binding the of virus to cells, while the A33 protein and B5 protein on EEV are considered to mediate the spread of virus between cells. Polyclonal and monoclonal antibodies against these four proteins have high level neutralizing activity to vaccinia virus (Gilchuk et al., 2016; Kaever et al., 2014; Law and Smith, 2001; Lustig et al., 2005; Paran and Lustig, 2010; Zajonc, 2017), monkeypox virus (Gilchuk et al., 2016), cowpox virus (Gilchuk et al., 2016) and variola virus (Gilchuk et al., 2016). Recombinant protein platform and DNA platform used to deliver A27, L1, A33, B5 alone or in combination can induce protective neutralizing antibody responses in mice (Hooper et al., 2000; Hooper et al., 2003; Hooper et al., 2004; Reeman et al., 2017) and monkeys (Buchman et al., 2010) against vaccinia virus and monkeypox virus with no safety concern observed. Sequence analysis confirm that these four vaccinia viral antigens and their homologous proteins in monkeypox virus (A29, M1, A35 and B6, respectively) and smallpox virus are highly conserved (the conservation score of all antigens are ≥93%, among which L1 is the most conserved, with a conservation ≥98.8%) (Hooper et al., 2003). These suggest that A27, A33, B5, and L1 can be used as targets for the development of vaccines against vaccinia and potentially other orthopoviruses such as monkeypox. In this study, monkeypox virus antigens were chosen for testing such vaccine’s cross reactivity.

At present, the available pox vaccines include the following: attenuated vaccinia virus ACAM2000 by Acambis (2^nd^ generation) and further attenuated (replication-defective in mammalian cells) vaccinia virus (Ankara) vaccine MVA-BN or Jynneos (3^rd^ generation) by Bavarian Nordic and a similar LC16m8 by Chiba (with restricted use in Japen). Although, these vaccines have been proved efficacious against smallpox in clinical trials (Kennedy and Greenberg, 2009a; Kenner et al., 2006a; Nalca and Zumbrun, 2010) and approved for containment of the recent outbreak of monkeypox, the vaccines face problems such as side effects, weak immunogenicity and low productivity to meet market need. New vaccines with higher efficacy and productivity as well as better safety profile are in urgent demand. Recently, mRNA vaccines have attracted much more attention due to its excellent immunogenicity, short preparation time and high yield and good safety data, which have been verified by intensive use of the two marketed COVID19 mRNA vaccines.

In this study, we have developed a mRNA vaccine candidate expressing four vaccinia viral antigens A27, L1, A33 and B5 in tandem in one molecule delivered by a LNP system to evaluate the immunogenicity against vaccinia and cross immunogenicity against other poxviruses in a mouse model. Immunization of mice with the candidate mRNA vaccine induced excellent vaccinia antigen-specific binding antibodies, neutralizing antibody responses and cellular immune responses. Strikingly, the sera from the vaccine-immunized mice cross reacted with all four monkeypox homologous antigens, holding promise for this mRNA vaccine candidate to be used for protection of broad orthopoxvirus infection.

## Results

### Design and synthesis of the mRNA molecule

A conventional mRNA sequence was designed to contain a cap structure, 5’UTR untranslated region, CDS translated region, 3’UTR untranslated region and a PolyA structure. The CDS translation region expresses four tandem VACV antigen molecules (A27, L1, A33, B5) separated by linkers, and the mRNA molecule was named mRNA-ALAB. A T7 promoter was added to the 5’ end of the complete mRNA sequence and a BspQI restriction enzyme cutting site was added to the 3’ end followed by cloning of the molecule into the synthetic plasmid vector pUCYH (Figure 1A). Digestion of the plasmid with BspQI can effectively linearize the plasmid template (Figure 1B). The mRNA-ALAB molecule was synthesized by in vitro transcription reaction using linearized plasmid as templates. Good integrity (Figure 1C) and purity of the synthesized mRNA (Figure 1D) were demonstrated using agarose gel electrophoresis and HPLC-SEC analysis. The capping efficiency of the mRNA-ALAB molecule was higher than 98% (data not shown).

**Figure 1.**
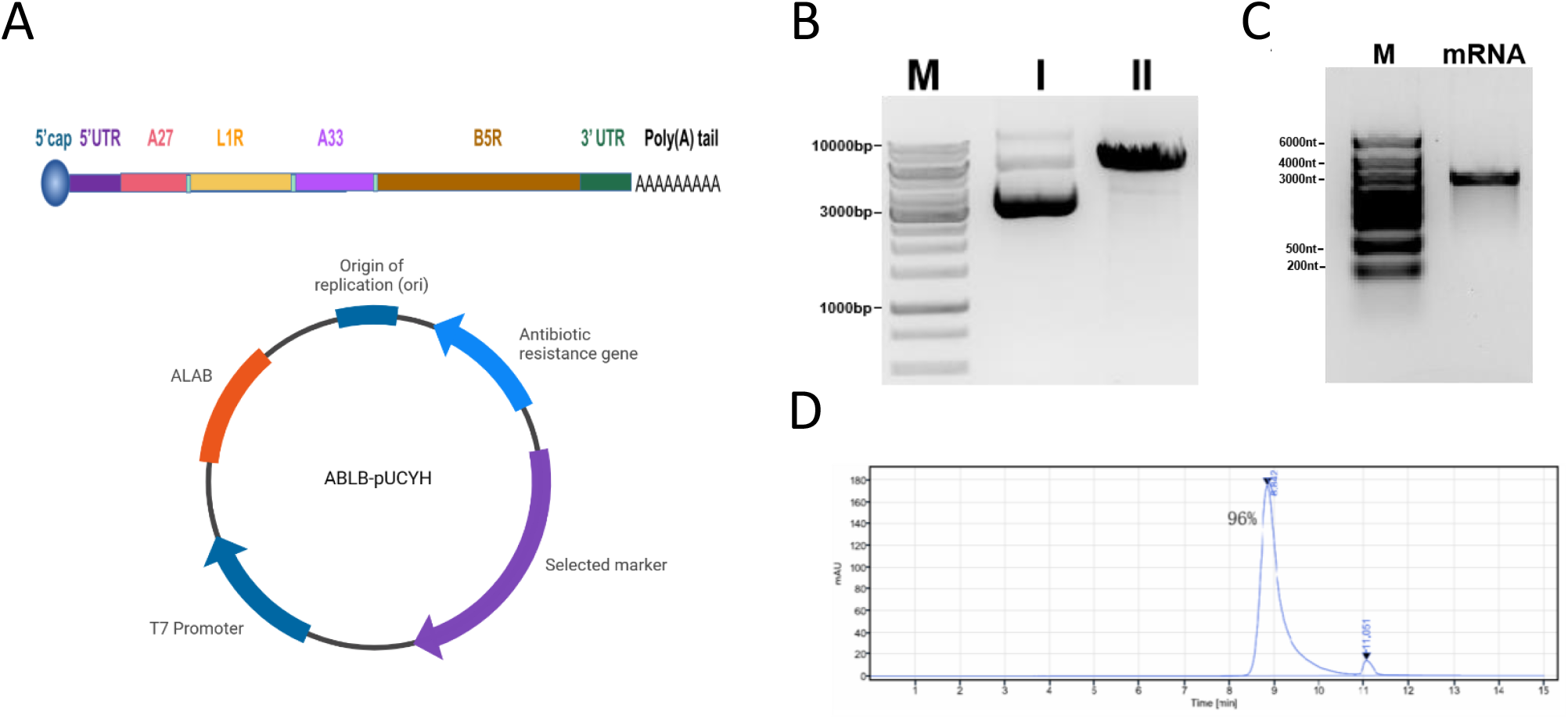
mRNA-ALAB structure design and preparation. (A) Structural diagram of the mRNA designed (up) and recombinant plasmid for the ALAB-DNA (down). (B) Agarose gel electrophoresis of BspQI enzyme digestion of ALAB-pUCYH plasmid. M, DNA marker. I, the supercoiled plasmid. II, Linearized plasmid after digestion. (C) Agarose gel electrophoresis of purified mRNA. (D) Detection of purified mRNA by HPLC-SEC analysis.

### Preparation and characterization of mRNA-ALAB-LNP Prototype

In this research, the mRNA-ALAB and lipids were fully mixed using microfluidic technology and formed mRNA-ALAB-LNP lipid nanocomplex (Figure 2A). Particle size of the mRNA-ALAB-LNP vaccine prototype was about 84 nanometers, while the empty LNP was about 75 nanometers (Figure 2B). The polymer dispersity index (PDI) of the mRNA-ALAB-LNP vaccine prototype and the empty LNP were all less than 0.1 (Table 1), indicating good uniformity of the nanoparticles. The encapsulation efficiency (EE) of mRNA in LNP was approximately 98% (Table 1). The transfection efficiency of mRNA-ALAB-LNP was demonstrated by flow cytometry data showing that transfected HEK293T cells were 20%, 85%, 6% and 75% positive for anti-A27, anti-L1, anti-A33 and anti-B5 staining, respectively (Figure 3, Figure S1).

**Figure 2.**
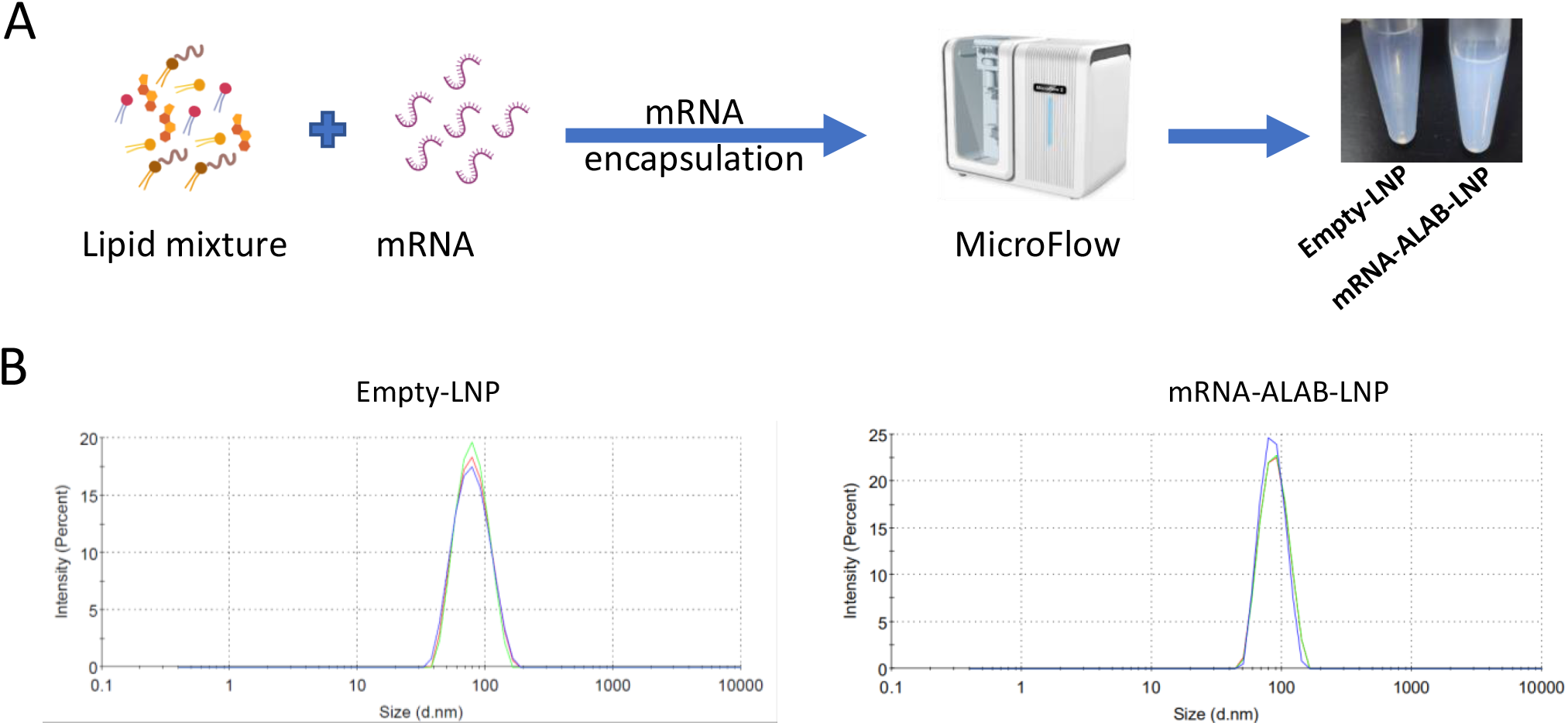
Preparation of mRNA vaccine candidate mRNA-ALAB-LNP. (A) Schematic diagram of the mRNA-ALAB-LNP vaccine preparation. (B) Particle size of mRNA-LNP.

**Table 1.** Characterization of mRNA-ALAB-LNP and Empty-LNP.

| Formulation | Size(nm) | PDI | EE |
| --- | --- | --- | --- |
| Empty-LNP | 75.7 | 0.07 | - |
|  | 75.9 | 0.05 |  |
|  | 74.2 | 0.09 |  |
| ALAB-LNP | 84.6 | 0.04 | 98% |
|  | 85.6 | 0.01 |  |
|  | 83.2 | 0.01 |  |

**Figure 3.**
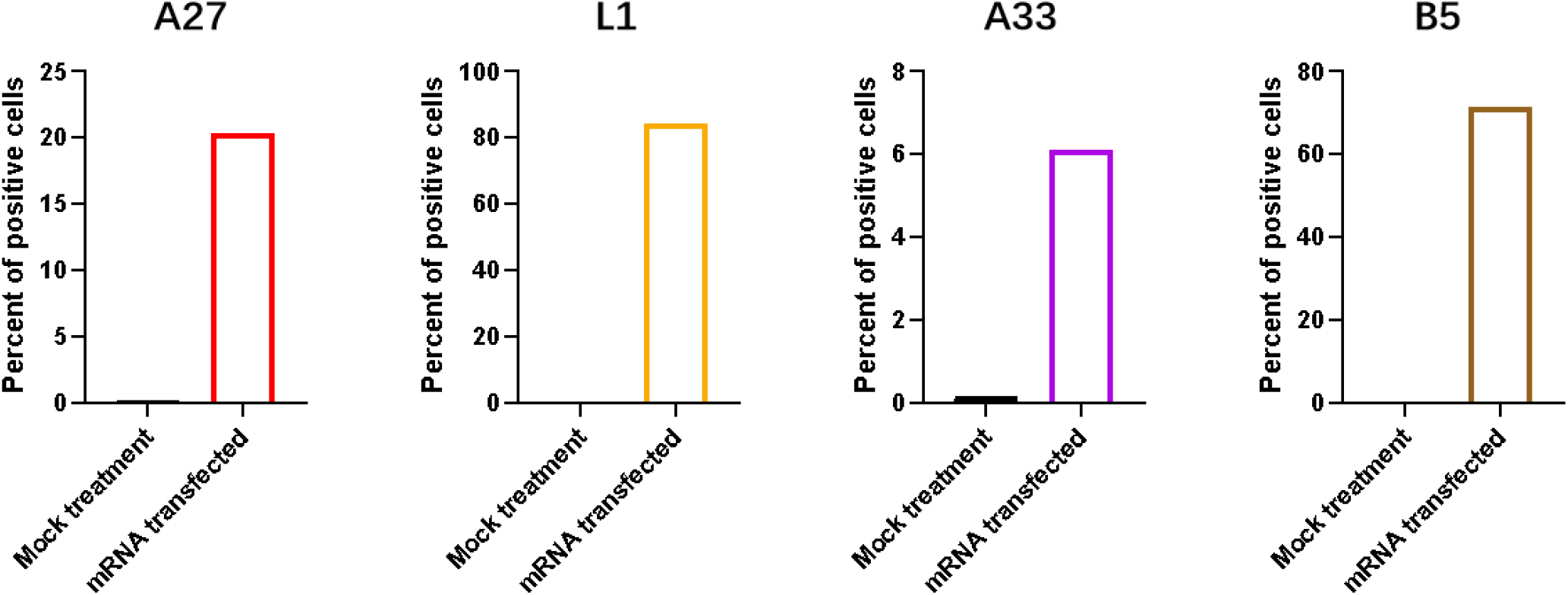
Expression of all four target proteins after mRNA-ALAB-LNP transfection. (A-D) Percentage of antigen-expressing cells was detected by flow cytometry. Expression of target proteins (A27, L1, A33, B5) in mRNA-ALAB-LNP transfected cells were determined by immune staining with anti-A27-, anti-L1-, anti-A33-, anti-B5-serum respectively, using mock treated cells as negative control.

### Prototype mRNA-ALAB-LNP vaccine induced excellent antigen-specific binding antibody responses in mice

In order to study the immunogenicity of the mRNA vaccine, 6-8 weeks old BALB/c mice were immunized by intramuscular injection twice at a 2-week interval (Figure 4A) with a dose of 20ug per injection. Blood samples were collected on day -3 (before prime), day 14 (2 weeks post prime), day 28 (4 weeks post prime), and day 42 (2 weeks post boost) for detection of antigen-specific binding antibodies (Figure 4B-4E). Spleens were taken at 45 days after second immunization for evaluation of cellular immunity. Strong anti-L1-specific and moderate anti-A33-, anti-A27- and anti-B5-specific antibody responses were induced after the first immunization and the antibody responses against all four antigens increased overtime. The mean titers of anti-A27 antibodies increased from 2,040 at 2 weeks post prime to 3,000 at 4 weeks post prime. Similarly, the anti-L1 antibodies increased from 24,840 (2 weeks post prime) to 55,080 (4 weeks post prime), anti-A33 antibodies increased from 2,680 (2 weeks post prime) to 17,640 (4 weeks post prime), and anti-B5 antibodies increased from 360 (2 weeks post prime) to 1,320 (4 weeks post prime). The antibody levels against the four antigens increased significantly after the boost immunization, with mean antibody titers of anti-A27, anti-L1, anti-A33, and anti-B5 reaching 110,000, 810,000, 14,580000 and 202,000, respectively. These results proved that the mRNA-ALAB-LNP vaccine candidate has induced potent VACV antigen specific binding antibody responses (Figure 4B-4E).

**Figure 4.**
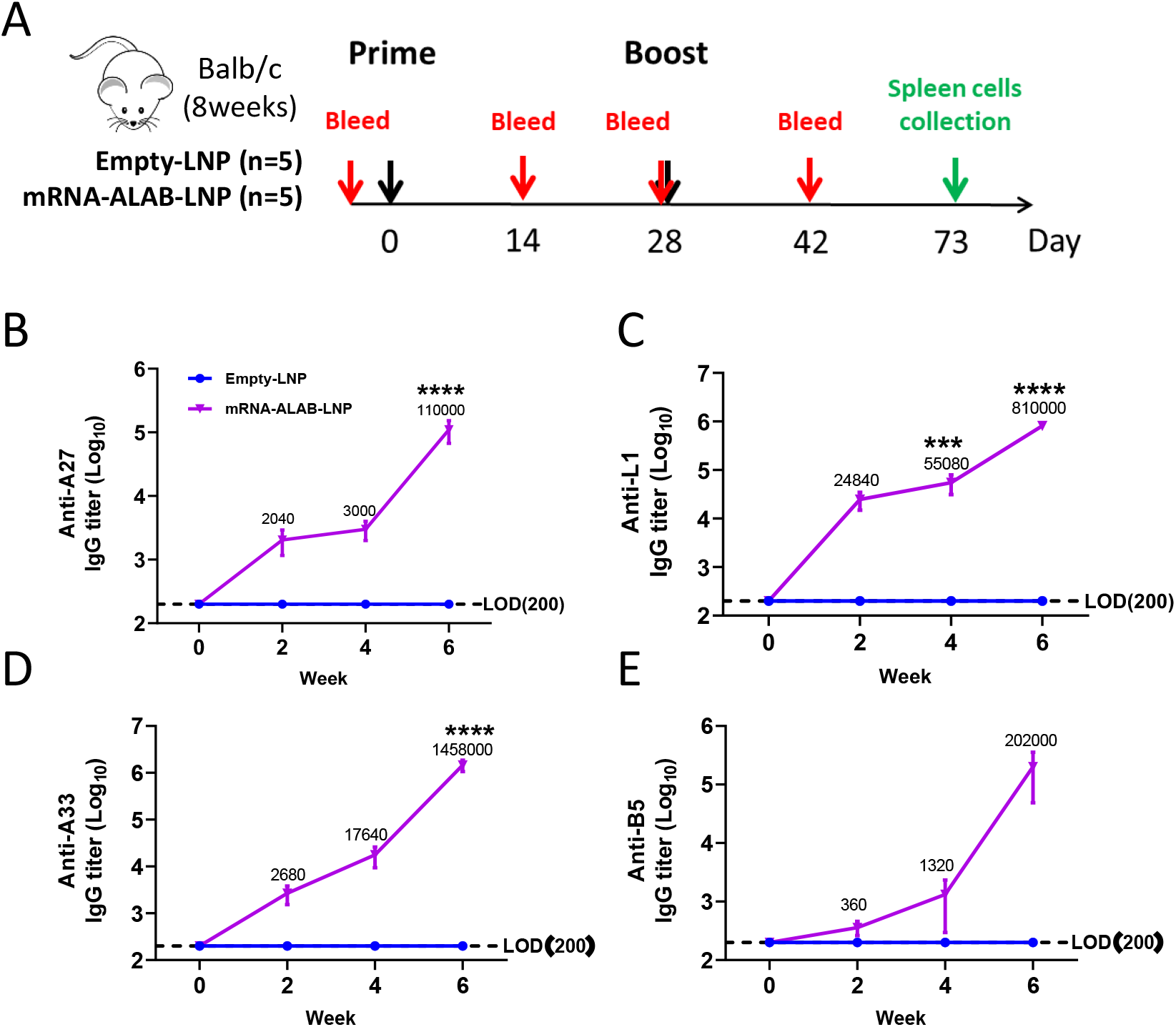
mRNA-ALAB-LNP elicits potent antibody response in mice. (A) Mice immunization schedule (n=5 for each group). The black and red arrows represent time points for immunizations and blood collection. (B-E) Serum binding antibody titer against A27 (B), L1 (C), A33 (D) and B5 (E) at different time points were tested. Statistical significance was assessed by two-way ANOVA with Sidak’s multiple comparisons test. Data are shown as means ± SEM.

### mRNA-ALAB-LNP vaccine immune sera showed potent poxvirus neutralization activity

In order to determine whether the antibodies produced by the mRNA-ALAB-LNP vaccine have virus neutralization activity, we incubated the inactivated serum from two weeks after boosting with 100 CCID50 of VACV virus for detection of neutralizing activity. Neutralizing activity of the serum was calculated by observing and grading the cytopathic effect (CPE). Both viral control (VC) and the immune serum from empty-LNP immunized mice, at a dilution of 1:80, did not exhibit neutralizing activity with obvious CPE shown in infected BSC-1 cells (Figure 5A). On the contrary, the immune serum from mRNA-ALAB-LNP vaccinated mice, at a dilution of 1:80 and 1:640 and even at 1:1280, showed protection of cells from virus infection with no CPE seen in infected BSC-1 cells (Figure 5A). The mean neutralizing antibody titer of mRNA-ALAB-LNP vaccinated group could reach 1,431 in this assay (Figure 5B).

**Figure 5.**
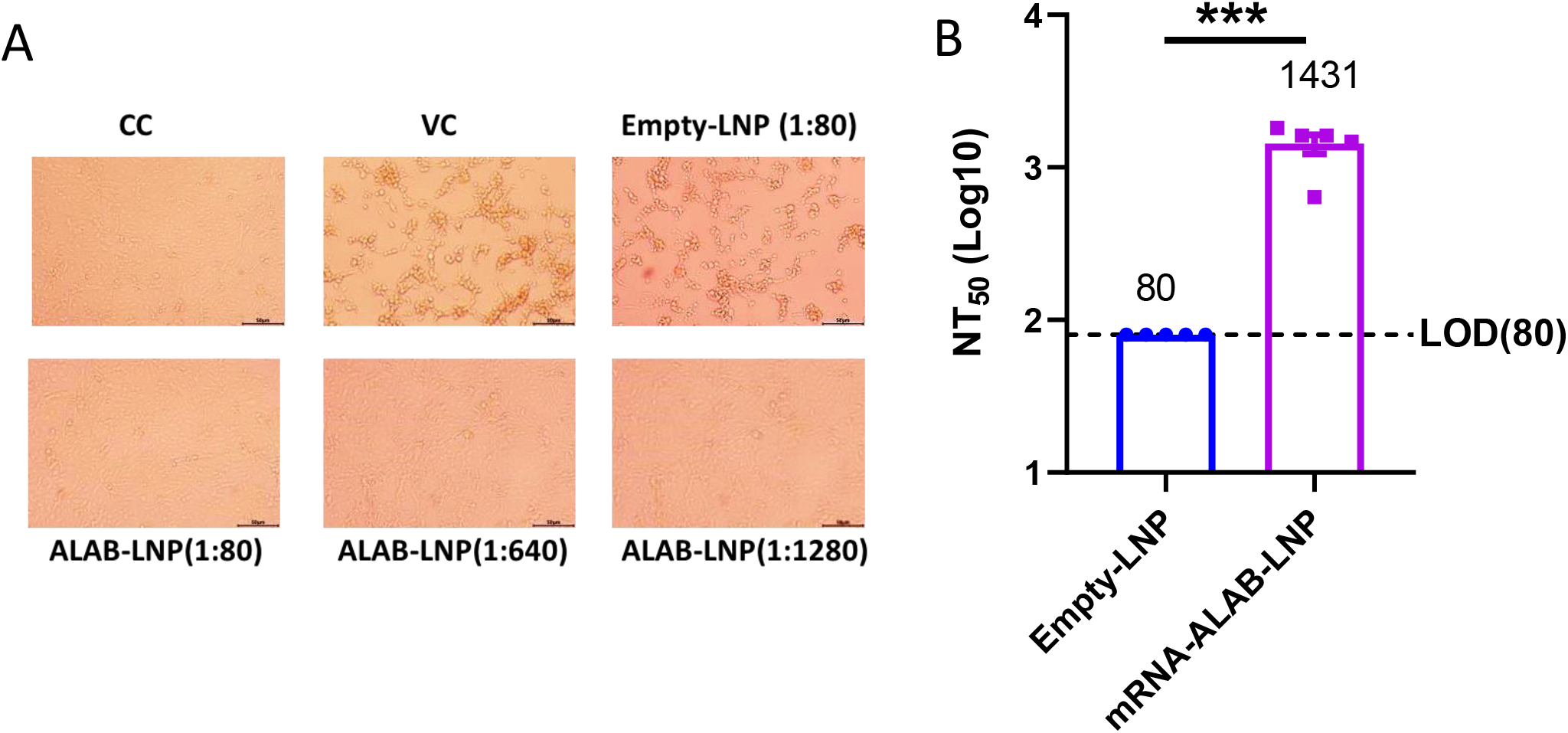
mRNA-ALAB-LNP elicits strong neutralizing antibody against vaccinia virus. (A) Representative images of VACV neutralization assay. Cell control (CC), viral control (VC); (B) VACV neutralizing antibody titers of immune serum (n=5). Statistical significance was assessed by two-tailed unpaired Student’s t-test. Data are shown as means ± SEM.

### mRNA-ALAB-LNP vaccine induced antigen specific cellular immune responses in mice

To explore whether the mRNA vaccine can induce specific cellular immune responses in mice, the mouse spleenocytes were collected at 45 days post boosting for detection of IFN-γ positive T cells specific to 6 antigen peptide libraries A27, L1-1, L1-2, A33, B5-1, B5-2, respectively using ELISPOT assay. Specific IFN-γ responses were detected to all four antigens and the responses were significantly higher than that of the control group immunized with empty LNP (Figure 6A). The A27 peptide pool and the A33 peptide pool stimulated higher level of IFN-γ with 904 spots/million cells and 2,747 spots/million cells recorded, respectively. In line with the highest total specific IgG response being anti-A33 (Figure 3D), the highest IFN-γ response was also seen in response to A33 peptide pool (Figure 6B). Moreover, no cellular immunity was detected against the linker.

**Figure 6.**
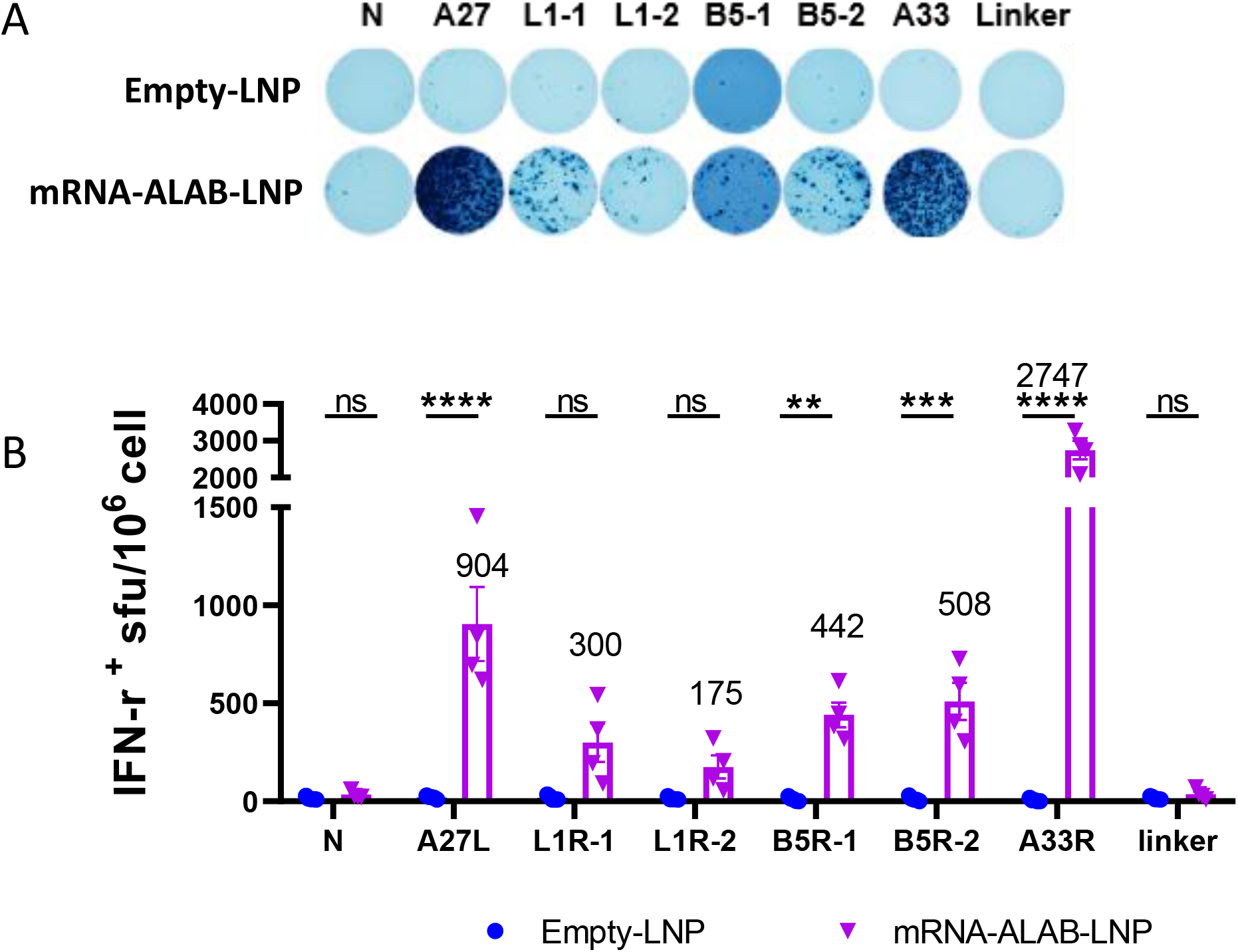
mRNA**-**ALAB-LNP elicits cellular immune response in mice (n=4). (A) Representative IFN-γ+ T cell immunospots data from mice spleenocytes stimulated with peptides. (B) Summary data of IFN-γ+ -secreting T cell numbers in spleenocytes post stimulation. Statistical significance was assessed by two-way ANOVA with Sidak’s multiple comparisons test. Data are shown as means ± SEM.

### Sera from mRNA-ALAB-LNP vaccinated mice showed high level cross-binding activity to monkeypox virus homologous antigens

Previous studies have shown that the four vaccinia virus antigens (A27, L1, A33, B5) are highly conserved among monkeypox virus, cowpox virus, and smallpox virus (Hooper et al., 2003). In this study, we tested whether the mRNA vaccine encoding these antigens could generate cross immunity against monkeypox homologous proteins A29, M1, A35 and B6, respectively. The immune sera from mice immunized with mRNA-ALAB-LNP were serially diluted and incubated with four proteins A29, M1, A35, and B6 to detect cross-binding antibody titers. ELISA results showed that strong serum IgG responses were seen in response to respective monkeypox antigens A35, M1, A29 and B6 after the first immunization and the titers increased overtime. (Figure 7A-7D). The average titers of anti-A29 antibody increased from 2,040 (2 weeks post prime) to 4,440 (4 weeks post prime), anti-M1 antibody increased from 74,520 (2 weeks post prime) to 184,680 (4 weeks post prime), anti-A35 antibody increased from 2,520 (2 weeks post prime) to 9,000 (4 weeks post prime), and anti-B6 specific antibody increased from 5,880 (2 weeks post prime) to 10,920 (4 weeks post prime). Furthermore, all four antibody levels increased significantly 2 weeks after boosting immunization, with antibody titers of anti-A29, anti-M1, anti-A35 and anti-B6 reaching 110,000, 2,430,000, 558,000, and 1,782,000, respectively. Collectively, these results demonstrated that mRNA-ALAB-LNP vaccine encoding vaccinia antigens (A27, L1, A33 and B5) induced equivalent or better cross-reactive antibodies against respective monkeypox antigens (A29, M1, A35, and B6) in mice. (Figure 7A-7D).

**Figure 7.**
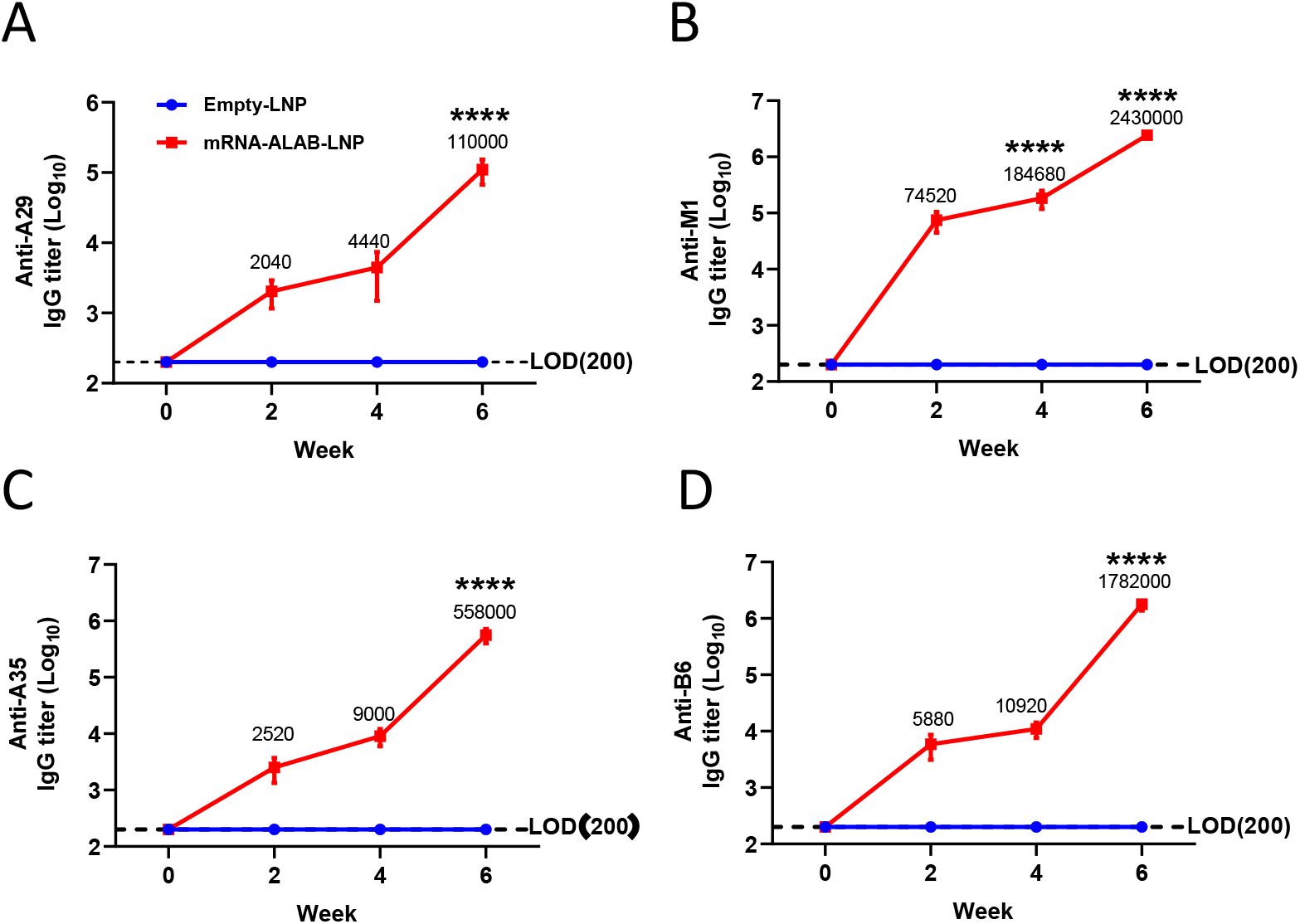
mRNA**-**ALAB-LNP elicits potent cross-reactive antibodies against MPXV antigens. (A-D) Serum binding antibody titer against A29 (A), M1 (B), A35 (C) and B6 (D) were tested at different time points. Statistical significance was assessed by two-way ANOVA with Sidak’s multiple comparisons test. Data are shown as means ± SEM.

## Discussion and Conclusion

Many efforts have been made to develop poxvirus vaccines, including live attenuated virus vaccine technology (Monath et al., 2004; Nalca and Zumbrun, 2010), replication-defective virus vaccine technology (Kennedy and Greenberg, 2009b; Kenner et al., 2006b), DNA vaccine technology (Heraud et al., 2006; Hooper et al., 2004), and recombinant protein technology (Buchman et al., 2010). Each has its own limitation such as severe skin side effects of live attenuated vaccine (Monath et al., 2004), poor immunogenicity of replication-deficient virus vaccines (Zaeck et al., 2023) and DNA vaccines (Heraud et al., 2006), and long development cycle of recombinant protein technology (Funk et al., 2021). Encouraged by the two new COVID19 mRNA vaccines, some researchers have also tried to use mRNA technology to develop monkeypox vaccines, mainly through design of multiple mRNAs to express a single or fused monkeypox antigen to induce antigen-specific humoral and cellular immunity (Alec et al., 2022; Fang et al., 2023; Hou et al., 2022; Sang et al., 2022; Zhang et al., 2023). Some of those design elicits potent immunogenicity, but will face challenges to deliver multiple mRNAs during process development and manufacturing. In this study, we designed a mRNA vaccine candidate to deliver four vaccinia antigens into cells in a single mRNA molecule. With regard to immunogenicity, this vaccine candidate induced strong humoral immune responses even after a single immunization in mice (yet to be tested in human). The efficiency of novel mRNA molecular design was also verified through induction of strong VACV-specific cellular immunity in mice. Furthermore, the potency of the four selected immune targets was proved by induction of significant levels of neutralizing antibody response. Yet, no obvious safety issue was observed after the mRNA vaccination in the mouse model, indicating a better safety. Additionally, the mRNA vaccine platform shortens the development cycle and the mRNA molecular design simplifies the manufacturing process, making the vaccine production process more feasible.

The purpose of this study is not only to generate a more efficacious and safer vaccine with a bigger productivity against potential crisis of smallpox, but also to develop a broader vaccine to provide protection against future pandemic that possibly caused by zoonotic poxviruses such as monkeypox, cowpox and camel pox, given that the recent monkeypox epidemic raised the poxvirus public health and safety concern and that cowpox and camelpox have been reported infectious to human. The sequence homology, morphological indistinguishability and antigenical relation are the basis for a broad poxvirus vaccine, pending on validation in animal studies and human trials. In this study, the quadrivalent mRNA-ALAB-LNP encoding four vaccinia viral antigens elicited equivalent or better binding antibody responses to monkeypox homologous antigens. The cross humoral reactivities between vaccinia and monkeypox were well demonstrated. The cross reactivity induced by the mRNA vaccine encoding vaccinia viral antigens to other poxviral antigens will be further studied. The vaccinia antigen-specific antibodies induced by the mRNA-ALAB-LNP vaccine have demonstrated potent neutralizing activity of vaccinia virus. The neutralizing capability of monkeypox and possibly other poxviruses by this vaccine will be evaluated in future studies.

In summary, we have developed a novel quadrivalent mRNA-ALAB-LNP vaccine candidate potentially against orthopoxviruses based on vaccinia viral antigen A27, L1, A35 and B5. The vaccine candidate elicited significant antibody response against vaccinia virus and monkeypox. The data demonstrated feasibility of mRNA-ALAB-LNP vaccine design and provided initial evidence of its potential as a broad orthopoxvirus vaccine candidate for its future optimization.

## Methods

### Ethic Statement

All experiments were performed strictly in accordance with the guidelines of care and use of laboratory animals by the Ministry of Science and Technology of the People’s Republic of China. Animal protocols were approved by the Animal Care and Use Committee of Yither Biotech Co., Ltd.

### Cell and virus

Hela S3 and BSC-1 cell lines were purchased from Procell Life Science & Technology Co., Ltd. (Wuhan, China) and cultured in Ham’s F-12K and MEM medium (Gibco) supplemented with 10% FBS (Gibco) and 1% antibiotics (Gibco), respectively. Vaccinia virus was purchased from ATCC and grown in Hela S3 cells. The virus titer was determined in BSC-1 cells, adjusted to half of the cell culture infective dose (CCID50).

### Mice study

Female 6-to 8-week-old BALB/c mice used for the experiments were grown under specific pathogen-free conditions at Bikai Laboratory Animal Co., LTD. Mice were randomly divided into two groups and immunized twice at an interval of 4 weeks with mRNA vaccine candidate mRNA-ALAB-LNP (20ug) or the same volume of empty-LNP. Serum was collected before immunization and every 2 weeks after the first immunization for antigen-specific antibody detection and heterologous antigen cross-reactive antibody examination. Spleens were harvested for cellular immunity evaluation 45 days after the second immunization.

### Construction of recombinant plasmid

The coding region expressing A27, L1, A33 and B5 proteins in tandem was designed and named as the ALAB region. The target molecule was obtained by adding the T7 promoter sequence and 5’ UTR sequence including Kozak sequence to the upstream of ALAB region and stop codon TGATAA, 3’UTR, PolyA and BspQI cleavage sites to the downstream of ALAB region. The target gene was next cloned into pUCYH plasmid by recombination and the recombinant plasmid ABLB-pUCYH sequence was confirmed by sequencing.

### ALAB mRNA preparation

The recombinant plasmid ABLB-pUCYH was extracted and linearized by restriction endonuclease BspQI cleavage followed by the plasmid recovery and purification. In vitro transcription reaction (50ul) was performed at 37°C for 3h after vortexing of the mixture containing linearized plasmid template, ATP (100mM), GTP (100mM), CTP (100mM), UTP (100mM), Cap analogue (100mM, Vazyme), T7 RNA polymerase (200U/ul, Vazyme), 10X T7 Reaction Solution (Vazyme), RNase enzyme inhibitor (40U/ul, Vazyme), pyrophosphatase (0.1U/ul, Vazyme) and RNase Free H2O (Invitrogen™,10977015). After the IVT, 170ul RNase-free H2O, 5ul DNaseI enzyme (NEB, M0303L) and 25ul 10X DNaseI enzyme reaction solution (NEB, M0303L) were added and incubated at 37°C for 20 min. Purified mRNA was obtained using OligoDT beads (Vazyme, N401). mRNA integrity and purity were analyzed by Agarose Gel and HPLC-SEC.

### mRNA vaccine preparation

Purified mRNA was diluted to 167ug/ml with PH4.0, 50 mM citric acid buffer to obtain aqueous solution. The organic phase solution was prepared by dissolving ionizable lipid, DSPC (AVT, S01005), cholesterol (Sigma-Aldrich, C8667), and PEG2000-lipid (Avanti Polar Lipids, 880150P) in ethanol. The aqueous and organic phase solutions were siphoned into the INano ™ L injector at a flow ratio of 3:1 with a total flow rate of 12ml/min for mRNA encapsulation. The mRNA-LNP complex was next rapidly diluted in PBS (pH7.4) and centrifuged at 3000rmp at 4°C for 10 min in 100kD ultrafiltration tube (Millipore). The mRNA-LNP complex was concentrated to 0.4mg/ml followed by filtration using 0.22um filter membrane to obtain mRNA-ALAB-LNP vaccine. Particle size and PDI of mRNA-ALAB-LNP vaccine were analyzed by Malvern particle size analyzer. The encapsulation rate of the mRNA vaccine was determined by RiboGreen (Invitrogen™, R11490) nucleic acid dye.

### Detection of antigen expression by flow cytometry

Briefly, HEK293T cells were transfected with 5ug mRNA-ALAB-LNP or equal volume of Opti-MEM. After 24h culturing, cells were detached from the plate surface with PBS containing 3% FBS, followed by staining with aqua fluorescent reactive dye (Invitrogen, L34966A, 1:1000) to determine alive versa dead. For surface staining, cells were stained with anti-A33-serum, anti-B5-serum, anti-L1-serum at 1:200 dilution, respectively. For intracellular staining, cells were fixed with IC Fixation Buffer (Invitrogen, 00-8222-49), permeabilized (Invitrogen, Permeabilization Buffer, 00-8333-56) and stained with a mouse serum that recognizes A27 (1:200 dilution) followed by adding anti-mouse AF488 (Invitrogen, A55058, 1:1000) as secondary antibody. After staining, cells were acquired on a FACS Attune NxT Acoustic focusing cytometer and data were analyzed using FlowJo 10.8.1.

### Antigen-specific binding antibody assay

ELISA was used to detect the titer of antigen-specific antibody IgG in mouse serum. Each antigen (A27, A33, L1 and B5) was diluted to 1ug/ml with coating buffer and incubated overnight at 4 ° C in a 96-well plate. The serum was serially diluted by 2-fold and was next placed into the plate coated with 5% milk at 37°C for 1h. After washing with PBST, the plate was incubated with HRP-conjugated goat anti-mouse IgG (1:4000) (Southern Biotech, Birmingham, AL, USA) and color reaction was developed by adding TMB (Invitrogen™, 002023) for 5 min followed by adding 50μL of ELISA terminating solution (New Cell&Molecular Biotech, E40500) to stop the reaction and the absorbance was measured at 450/620 nm using an ELISA plate reader (Thermo Fisher, Varioskan LUX). The mean OD (optical density) of sera from naïve mice was multiplied by 2.1 to define the positive cutoff point. The cross-binding activity of immunized mice sera with monkeypox virus antigen (A29, M1, A35 and B6) was detected by ELISA as described above. Results are presented as mean titers of antibodies in the sera.

### Neutralizing antibody assay

To detect neutralizing antibodies, mice sera collected at 2 weeks post boosting were tested as follows: serum was inactivated at 56°C for 30 min, followed by 10 serial dilution (2-fold) with serum-free MEM medium (Gibco) starting from 1: 40. 50ul serum was next mixed with an equal volume of VACV (100 CCID50) and incubated at 37°C for 2h. The mixture was added to the 96-well cell plates with BSC-1 cells laid one day before (1×10^4^ /well). After 3 days of incubation at 37°C, CPE and cell growth status were observed to determine the numbers of virus-infected wells and the infection rate at each dilution was calculated. Finally, the dilution that can reduce 50% virus infection was defined as neutralizing antibody titers.

### Cellular immune response assay

The ELISPOT plate (Millipore, MSIPS4W10) was filled with 35% ethanol at 50ul/well for 1min. The liquid was discarded and the plate was washed with sterile deionized water at 200ul/well for 5 times. IFN-γ capturing antibody (MABTECH,3321-2H) was diluted to 15ug/ml with PBS and added to the above plates at 100ul/well and the plates were incubated at 4°C overnight. On the second day, the plates were washed with PBS, and RPMI 1640 medium (containing 10%FBS) was added followed by incubation at room temperature for 30min. Single cell suspension was obtained from a mouse spleen and cell numbers were counted after erythrocyte lysis. The culture medium in the plate was discarded and appropriate numbers of cells were added. Next, polypeptides were added at a final concentration of 2.5ug/ml each. The plates were placed in an incubator set at 37°C for 36h followed by discarding the cells and washing with PBS. The diluted detection antibody (1:1000) (MABTECH,3321-2H) was added and the plate was incubated at room temperature for 2h. The diluted streptavidin-HRP (1:1000) (MABTECH,3321-2H) was added and the plate was incubated at room temperature for 1h. Finally, TMB solution (MABTECH, 3651-10) was added followed by washing with deionized water after spots became obvious. After the plates were dried at room temperature, pictures were taken and the spots in the wells were counted using enzyme-linked immunospot analyzer (CTL, S6 Universal).

### Statistical analysis

Statistical analysis was done using GraphPad Prism 8.0 (GraphPad Software, San Diego, CA, USA) and results are presented as mean ± standard error of the mean (SEM). The exact sample size (n) for each experimental group is indicated in the figure legends. Differences between two groups were analyzed with unpaired t-tests or two-way analysis of variance (ANOVA). P values < 0.05 were considered statistically significant.

## Conflicts of Interests

All authors are employees of Yither Biotech and Ab&B Biotech. A patent has been filed related to this study.

## Acknowledgement

We would like to thank other colleagues involved in this work; This work was funded by Yither Biotech.

## Authors Contribution

Supervision, Z. L; Conceptualization, Y. X and Z. L; Methodology, C. S, Q Y and Z. L; Formal analysis, C.S and Z. L; Investigation, C. S, X. G, Y. W and Z. L; Writing Original Draft, Z. L; Editing and Review, C. S, C. Y, Y. X and Z. L; Visualization, C.S and Z. L;

## Figure Legends

**Figure S1.**
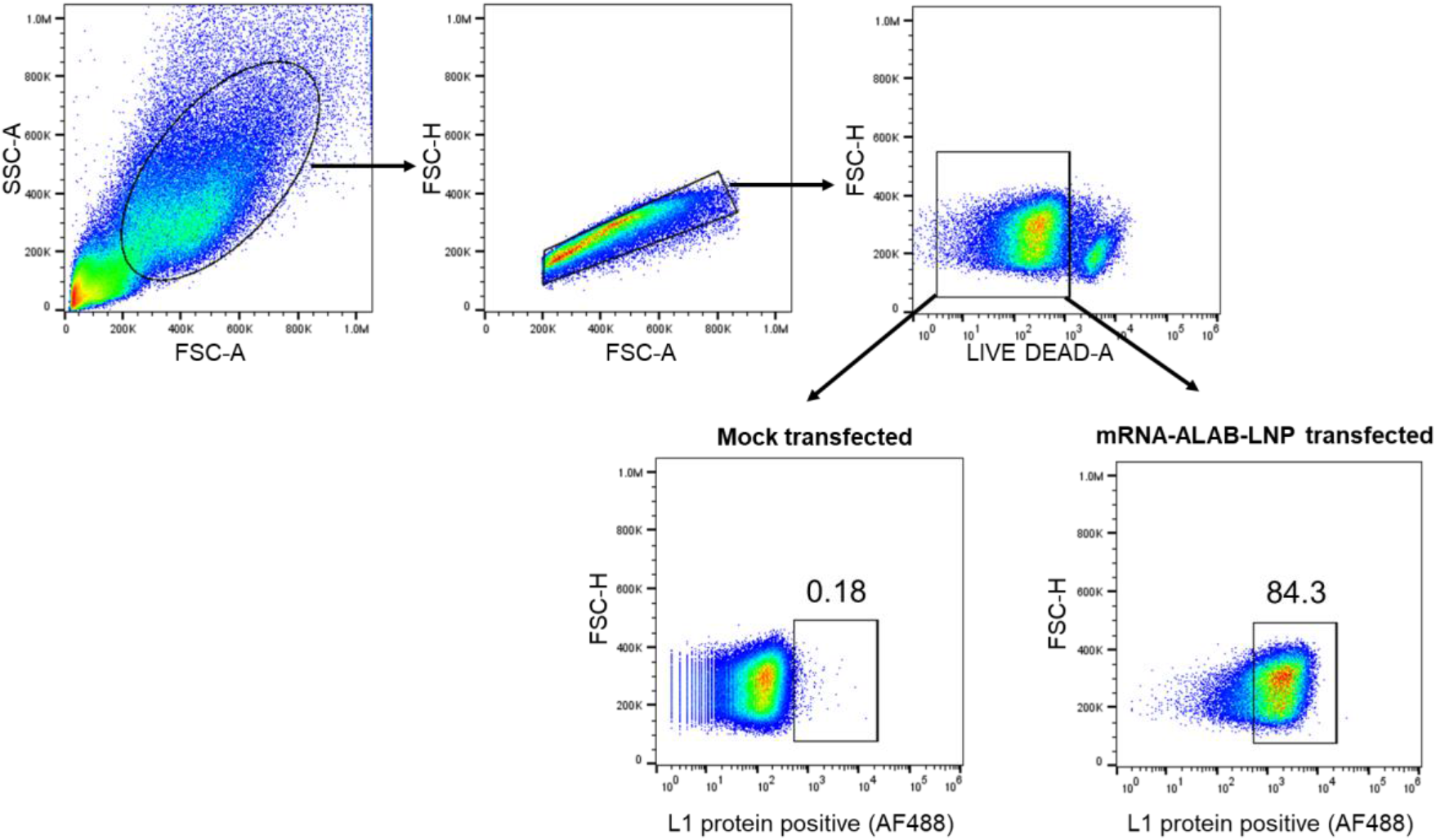
Representative gating strategy of flow cytometry. Mock-transfected and mRNA-ALAB-LNP transfected HEK293T cells were detected for expression of L1 protein.

## Notes

### Competing Interest Statement

The authors have declared no competing interest.

